# Genetic diversity analysis of main agronomic traits and ISSR markers in 35 ornamental rape germplasm resources

**DOI:** 10.1101/2024.07.29.605676

**Authors:** Mang Xia, Meizhu Chen, Xiaoxiao Dong, Jingdong Chen, Miao Cheng, Heping Wan, Yuanhuo Dong, Changli Zeng, Xigang Dai

**Affiliations:** Hubei Engineering Research Center for Protection and Utilization of Special Biological Resources in Hanjiang River Basin, College of Life Sciences, Jianghan University, Wuhan 430056, China

## Abstract

Rape (*Brassica napus* L.) is a major oil crop in our country, valued for its oil and ornamental uses. This study analyzed 35 ornamental rape germplasm resources from different origins to examine differences in agronomic traits and molecular markers. Nine agronomic traits were assessed in the field for variability, correlation, principal component analysis, and cluster analysis. Genetic diversity was analyzed using microsatellite (ISSR) markers and the unweighted pair group method with arithmetic mean (UPGMA). Our findings revealed a notable average coefficient of variation of 22.59% across the nine agronomic traits, with flower color exhibiting the highest variability and corolla width the least. The observed range of variation spanned from 9.24% to 83.38%, the correlation among these traits was generally low, with a mere 13.9% demonstrating significant correlations. The four principal components accounted for an impressive 84.62% of the cumulative contribution rate, while the genetic similarity, as gauged by eight ISSR primers, varied from 0.675 to 0.980. Most strikingly, we observed that plants from the same geographical region displayed molecular-level differences, underscoring the rich genetic diversity inherent in the 35 ornamental rape resources under study. Employing UPGMA cluster analysis on the primary agronomic traits and ISSR molecular markers, the 35 ornamental rape resources were categorized into seven and four distinct groups, respectively. Although the clustering outcomes from these two methodologies did not align perfectly, they served to complement each other. Collectively, these insights offer a theoretical framework for the innovation of ornamental rape germplasm resources and the cultivation of novel varieties.

## Introduction

Rape (*Brassica napus* L.), a member of the Brassica genus, is widely cultivated around the world. In our country, the annual planting area is approximately 66.67 million hectares, accounting for about one-third of the global yield and area [1-3]. Besides oil production, rape has various uses, including ornamental, vegetable, forage, and green manure applications. It plays a significant role in environmental protection, the development of eco-agriculture, and leisure agriculture [4-9].Traditionally, rape flowers are predominantly golden yellow. While large fields of golden rape flowers are visually striking, the uniform color can lead to aesthetic fatigue among tourists, thereby limiting the sustainable development of rape flower tourism. In contrast, ornamental rape varieties with diverse flower colors, abundant blooms, rich fragrance, and dual-purpose for sightseeing and oil production offer significant ornamental, tourism, economic, and social value. These varieties can be integrated into agriculture, rural development, tourism, and the catering industry, making them an ideal choice for promoting rural tourism.Therefore, studying the genetic diversity and differences among colored rape germplasm resources is of great significance. Such research can help identify and utilize superior germplasm resources, create new materials for ornamental rape breeding, and develop breakthrough varieties. This is crucial for enhancing the ornamental appeal and economic potential of rape flowers, thereby supporting the sustainable development of rural tourism.

Rape is easy to grow and manage, featuring bright colors, wide adaptability, and extensive planting areas. It is an ideal resource for leisure agriculture and flower sea tourism. The famous rape flower seas in China, such as those in Wuyuan County, Menyuan, Luoping, Hanzhong, and Xinghua, attract numerous tourists from both home and abroad for flower viewing, photography, and leisure activities, forming a unique tourism industry [10]. Ornamental rape is a new variety that differs from common rape in color and shape [11]. Artificial cultivation of ornamental rape began in the 1980s, and after a series of studies, improved varieties were first planted in Wuyuan and Luoping.With the advancement of rapeseed breeding techniques [12-18], it is now possible to breed various rape varieties with different colors and extended blooming periods, while maintaining their growth habits and oil quality [19-23]. This has significantly enhanced the ornamental and economic value of rape flowers, making them a key component in the development of rural tourism and leisure agriculture.

Germplasm resources, also known as gene resources, are the material basis for the improved gene sources of crop varieties and agricultural innovations. Studying the genetic diversity of ornamental rape germplasm resources is crucial not only for the collection and preservation of these resources [24], but also for analyzing their genetic relationships and backgrounds. Additionally, such studies can effectively evaluate and utilize ornamental rape resources, providing excellent materials for improving breeding quality and efficiency.In this study, 35 ornamental rape germplasm resources were analyzed to assess their genetic diversity based on major agronomic traits and ISSR molecular markers. The aim was to clarify the genetic background and relationships among these ornamental rape resources, and to identify superior ornamental rape resources. This research provides theoretical guidance for further innovation in ornamental rape germplasm resources and the utilization of heterosis.

## 2. Materials and Methods

### 2.1. Plant Material

Ornamental rape germplasm resources were provided by the Hannan base of Jianghan University. In 2022, the plants were cultivated in the greenhouses at the Hannan base, located in Caidian District, Wuhan, Hubei Province. The flowering period began on May 8, 2023, and the harvest took place on September 7, 2023. Wuhan has a tropical monsoon climate with abundant rainfall and an average annual temperature of 16°C. The annual precipitation is approximately 1300 mm, and the total number of sunshine hours ranges from 1810 to 2100 hours. These conditions are highly suitable for the growth of rape.

### 2.2. Measuring items and methods

#### 2.2.1. Investigation of agronomic traits

The nine agronomic characteristics of the aboveground part of rape during both vegetative and reproductive growth stages were investigated and analyzed. Three rape plants of the same variety were randomly selected for the measurement of agronomic traits. These traits were measured according to the standards of agronomic morphological measurement outlined in Table 1. The quality traits of ornamental rape were assigned based on the criteria specified in Table 2.

**Table 1.** Determination indexes of agronomic traits of Brassica napus.

| No. | Range | Agronomic Trait | Determination Method | Unit |
| --- | --- | --- | --- | --- |
| 1 | Vegetative growth | Plant height | Distance from rhizome bulge to plant apex | cm |
| 2 |  | Leaf length | Leaf length of mature middle leaves | cm |
| 3 |  | Leaf width | Leaf width of mature middle leaves | cm |
| 4 |  | Number of nodes in main stem | Number of stems scattered from the main stem | / |
| 5 |  | Leaf margin | Leaf margin type of mature middle leaves | / |
| 6 | Reproductive growth | Corolla width | Petals in full bloom in diameter | cm |
| 7 |  | Silique length | Leaf length of mature Silique | cm |
| 8 |  | 1000-seed weight | Clean the weight of 1000 seeds | g |
| 9 |  | Color of petals | Observe petal color in full bloom | / |

**Table 2.** Evaluation of quality traits in ornamental rape.

| **Trait** | **Assignment** |
| --- | --- |
| Leaf margin | 1:Round leaf; 2:wavy leaf;3:toothed leaf;4:lobed leaf |
| Color of petals | 1=Yellow;2=White;3=Red;4=Purple;5=Orange yellow;6=Pale yellow;7=Peach;8=Pinkish purple;9=Golden yellow;10=Apricot yellow;11=Carmine |

### 2.3. Isolation of genomic DNA

Fresh and young leaves was collected from 20 days old seedlings and total genomic DNA was extracted following the Cetyl Trimethyl Ammonium Bromide (CTAB) method with slight modifications. For each of the 35 ornamental rape samples, 1 g of young leaves was taken and placed in a mortar. Liquid nitrogen was added to make the leaves brittle, and they were then ground into a powder. The powder was transferred into numbered centrifuge tubes. To each tube, 1 mL of CTAB extraction solution and 20 μL of β-mercaptoethanol were added, mixed thoroughly, and then incubated in a 65°C water bath for one hour. After incubation, the samples were centrifuged at 12,000 rpm for 10 minutes using an IKA G-L centrifuge. The supernatant was transferred to new centrifuge tubes, and chloroform-isoamyl alcohol (24:1) was added in proportion. The mixture was centrifuged again at 12,000 rpm for 10 min. The supernatant was collected, and 500 μL of isopropanol was added, mixed by inverting the tubes for 5 min, and then placed in a -20°C freezer for 30 min to precipitate the flocculent DNA. The samples were centrifuged at 12,000 rpm for 10 minutes, the supernatant was discarded, and the pellet was washed with 1mL of 70% ethanol. The tubes were gently shaken for a few minutes to remove the ethanol, resulting in crude DNA. After air drying, the DNA was dissolved in 30 μL of ddH_2_O and fully dissolved for further use. The isolated genomic DNA was quantified using spectrophotometer (NanoDrop ND 1000) and quality was checked by electrophoresis on 1% (w/v) agarose gel. Finally, the DNA was diluted to 30 ng/μL for PCR amplification.

### 2.4. ISSR-PCR Amplification of ornamental rape

The ISSR primers were designed with reference to the ISSR primer sequences published by Columbia University (New York, NY, USA) and synthesized by Wuhan Qingke Biotechnology Co., Ltd. (Wuhan, China). A total of 8 ISSR primers were meticulously chosen from a pool of 30, based on their distinct, clearly resolved, and highly polymorphic amplified products (Table 3). Utilizing these primers, the DNA from 35 ornamental rape samples was amplified through the ISSR-PCR technique. The reaction system was composed of 2 μL of template DNA, 0.5 μL of primer, 11 μL of 2×Es Taq Master Mix (Dye), and ddH_2_O to reach a total volume of 20 μL. The PCR amplification protocol included an initial pre-denaturation step at 94°C for 5 min, followed by 40 cycles of denaturation at 94°C for 30 s, annealing at 52.2°C for 30 s, and extension at 72°C for 45 s, concluding with a final extension at 72°C for 5 min. The resulting amplified products were then subjected to electrophoresis on a 1% agarose gel, stained with ethidium bromide (EB), and subsequently visualized using a gel imaging system.

**Table 3.**
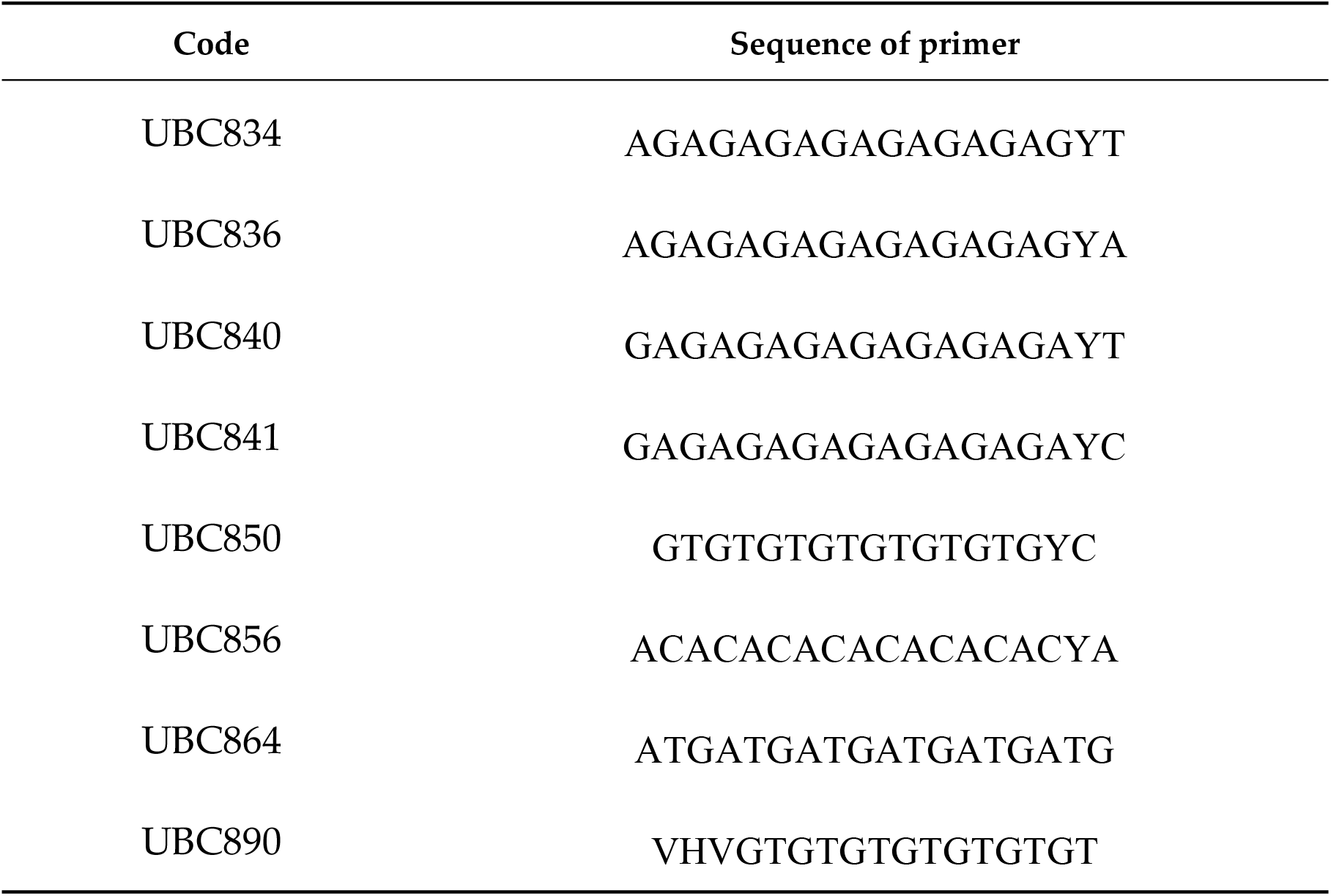
Details of 8 primers used in ISSR-PCR.

### 2.5. Statistical Analysis

Data were statistically analyzed using Excel 2019 and SPSS 19.0. Excel 2019 was utilized to compute and analyze the agronomic trait data, calculating the mean, maximum, minimum, standard deviation, and coefficient of variation. SPSS 19.0 was employed for correlation analysis and principal component analysis of the agronomic traits. The ISSR band was scored as binary data with presence of band indicated by 1 and its absence by 0. Data were statistically analyzed using the NTSYS-pc version 2.1 software.Cluster analysis was conducted using the Unweighted Pair Group Method with Arithmetic Mean (UPGMA), and a dendrogram illustrating the genetic relationships was constructed.

## Conclusion

### 3.1. Genetic Diversity Analysis of Ornamental Rapeseed Germplasm Resource

Significant variations were observed across nine agronomic traits among different germplasm sources of ornamental rape (Table S1). Within the 35 rape varieties assessed, the plant height varied notably, with variety No. 25 being the tallest at 54 cm, and the shortest being No. 20, reaching only 29.3 cm. Varieties No. 13, No. 21, and No. 31 were distinguished by their longer leaves, measuring 55.7 cm, 55 cm, and 55 cm, respectively. Notable differences were also found in leaf width, with the smallest leaves belonging to varieties No. 12 and No. 17, which were a minimum of 18.3 cm wide, whereas variety No. 21 exhibited the widest leaves, at 31.17 cm—approximately 1.7 times wider than the smallest. During the reproductive growth phase, varieties No. 3 and No. 10 demonstrated exceptional performance, with the best corolla width and 1000-grain weight among the 36 varieties, at 2.8 to 2.9 cm and 4.5 to 4.8 g, respectively. Additionally, varieties No. 15 and No. 19 showed superior silique length.

According to Table 4, when ranked by the magnitude of variation coefficients for agronomic traits, the order from the highest to the lowest is as follows: petal color > leaf margin > 1000-seed weight > silique length > leaf width > number of nodes in the main stem > plant height > leaf length > corolla width. The overall average variation coefficient is 22.6%. The petal color exhibited the highest variation coefficient at 83%, followed by leaf shape and 1000-seed weight, with 24% and 21% respectively. The corolla width had the lowest variation coefficient at 9%. These findings underscore the rich genetic diversity present in the 35 ornamental rape germplasm varieties, particularly in relation to agronomic and key quality traits.

**Table 4.** Diversity analysis of agronomic traits of different ornamental rape varieties.

| Agronomic Trait | Max | Min | Range | Mean | SD | CV (%) |
| --- | --- | --- | --- | --- | --- | --- |
| Plant height | 54.00 | 29.33 | 24.67 | 42.55 | 5.13 | 12.06 |
| Leaf length | 55.67 | 37.67 | 18.00 | 48.58 | 4.67 | 9.61 |
| Leaf width | 31.17 | 18.33 | 12.83 | 23.29 | 3.03 | 13.01 |
| Number of nodes in main stem | 18.33 | 10.67 | 7.66 | 12.68 | 1.53 | 12.07 |
| Corolla width | 2.87 | 2.00 | 0.87 | 2.49 | 0.23 | 9.24 |
| Silique length | 11.83 | 5.50 | 6.33 | 8.43 | 1.59 | 18.86 |
| 1000-seed weight | 4.76 | 1.84 | 2.92 | 3.42 | 0.72 | 21.05 |
| Leaf margin | 4.00 | 1.00 | 3.00 | 2.66 | 0.64 | 24.06 |
| Color of petals | 11.00 | 1.00 | 10.00 | 3.31 | 2.76 | 83 |

### 3.2. Correlation analysis of agronomic traits in ornamental rapeseed germplasm resources

Table 5 presents the correlation analysis of nine agronomic traits, revealing that plant height is not only significantly and positively correlated with leaf length (P<0.01) but also shows a positive correlation with the count of main stem branches (P<0.05). This relationship extends to leaf length, which in itself is highly and positively correlated with both leaf width and silique length (p<0.01). Echoing this pattern, leaf width is found to have a highly significant positive correlation with silique length (P<0.01). Additionally, a strong positive correlation is observed between corolla width and the weight of 1000 seeds (P<0.01).

**Table 5.** Analysis results of correlation between agronomic traits of ornamental rapeseed.

| Agronomic Trait | Plant height | Leaf length | Leaf width | Number of nodes in main stem | Corolla width | Silique length | 1000-seed weight | Leaf margin | Color of petals |
| --- | --- | --- | --- | --- | --- | --- | --- | --- | --- |
| Plant height | 1.000 |  |  |  |  |  |  |  |  |
| Leaf length | 0.447** | 1.000 |  |  |  |  |  |  |  |
| Leaf width | 0.096 | 0.654** | 1.000 |  |  |  |  |  |  |
| Number of nodes in main stem | 0.340* | 0.261 | 0.291 | 1.000 |  |  |  |  |  |
| Corolla width | 0.297 | 0.169 | -0.057 | -0.047 | 1.000 |  |  |  |  |
| Silique length | 0.249 | 0.510** | 0.489** | 0.263 | 0.253 | 1.000 |  |  |  |
| 1000-seed weight | 0.134 | -0.073 | -0.01 | -0.077 | 0.602** | 0.095 | 1.000 |  |  |
| Leaf margin | -0.117 | -0.086 | 0.123 | -0.017 | -0.003 | 0.187 | 0.002 | 1.000 |  |
| Color of petals | -0.317 | -0.058 | -0.069 | -0.024 | 0.033 | -0.138 | -0.083 | 0.046 | 1.000 |

\*\* indicates extremely significant differences at the 1% level ( $P < 0.01$ ) and \* indicates significant differences at the 5% level ( $P < 0.05$ )

These insights shed light on the complex interplay among the agronomic traits in ornamental oilseed rape. Grasping these correlations is essential for informed breeding practices and the selection of cultivars, with the goal of producing varieties that exhibit both desirable growth habits and enhanced ornamental features. It is crucial, however, to recognize that the presence of these correlations does not automatically suggest a direct causal link. As such, further research is warranted to delve into the genetic and environmental factors at play, which may be shaping these trait relationships.

### 3.3. Principal component analysis of agronomic traits in ornamental rapeseed germplasm resource

When using principal component analysis (PCA) for agronomic traits, the suitability of the data for PCA was evaluated. As can be seen from Table 6, the test statistic KMO (Kaiser-meyer-olkin) is 0.534, which can be analyzed.

**Table 6.** KMO and Bartlett’s test results.

| KMO value | Bartlett spherical degree test |  |  |
| --- | --- | --- | --- |
|  | Approximate chi-square | df | P value |
| 0.534 | 69.558 | 21 | 0.000 |

As detailed in Table 7, principal component analysis (PCA) yielded four principal components with a combined contribution rate of 84.62%. This substantial cumulative contribution indicates that these four components effectively encapsulate the primary characteristics of the 35 ornamental rape germplasm resources under study.

**Table 7.** KMO and Bartlett’s test results.

| Agronomic Trait | Principal component |  |  |  | Total load | Best indicators ranking |
| --- | --- | --- | --- | --- | --- | --- |
|  | 1 | 2 | 3 | 4 |  |  |
| Plant height | -0.170 | -0.052 | 0.833 | 0.014 | 0.04 | 6 |
| Leaf length | 0.364 | -0.147 | 0.352 | -0.306 | 0.12 | 4 |
| Leaf width | 0.508 | -0.026 | -0.298 | 0.033 | 0.14 | 2 |
| Number of nodes in main stem | -0.095 | 0.029 | -0.077 | 1.007 | 0.06 | 5 |
| Corolla width | 0.002 | 0.476 | 0.165 | -0.139 | 0.12 | 4 |
| Silique length | 0.388 | 0.142 | -0.137 | 0.043 | 0.16 | 1 |
| 1000-seed weight | -0.003 | 0.615 | -0.257 | 0.166 | 0.13 | 3 |
| Eigenvalues | 2.55 | 1.68 | 0.99 | 0.71 | / | / |
| Contribution rate(%) | 36.42 | 24 | 14.09 | 10.12 | / | / |
| Cumulative contribution rate(%) | 36.42 | 60.42 | 74.50 | 84.62 | / | / |

The first principal component, accounting for the highest contribution at 36.42%, was predominantly influenced by traits such as leaf length, leaf width, corolla width, and silique length, all of which exhibited substantial loadings with positive values. The second principal component had a contribution of 24%, with corolla width and 1000-seed weight being the traits with significant positive loadings. Following this, the third principal component contributed 14.09% to the variance, where plant height, leaf width, and 1000-seed weight were the main traits with higher negative loadings. The fourth principal component added a further 10.12% to the overall explanation of variance.

To determine the impact of each trait on ornamental rape, the loadings of the seven phenotypic indices in the four principal components were multiplied by their respective variance contribution rates and then aggregated. The results indicated that, in descending order of impact, pod length, leaf width, 1000-grain weight, leaf length, corolla width, the number of branches on the main stem, and plant height are the key factors that successively influence the quality of Brassica napus.

### 3.4. Cluster analysis

Utilizing nine agronomic traits, the ornamental rape germplasm resources were categorized into four distinct groups (Figure 1). The first group comprises 9 varieties, which are characterized by a short plant stature, brief leaf length, narrow leaf width, limited main stem branching, and reduced corolla size and 1000-grain weight. The second group is made up of 6 varieties, distinguished by their elongated leaves, broad leaf span, and relatively narrow corolla width. The third category encompasses 5 varieties, marked by their impressive plant height, abundant main stem branches, expansive corolla, medium-length siliques, and notably heavy 1000-grain weight. The fourth group is the most diverse, with 15 varieties showcasing long and broad leaves, a generous number of main stem branches, large corollas, extended silique lengths, and substantial 1000-grain weight.

From these classifications, it is evident that the 5 ornamental rape accessions within the third category excel in agronomic traits when juxtaposed with the other 30 accessions. While their leaf length and width may not be the most superior, the accessions excel in other critical attributes of ornamental rape, such as the number of main stem branches, corolla width, and plant height, which are paramount for their ornamental value.

### 3.5. ISSR analysis

The eight selected ISSR primers were capable of amplifying distinct bands from 35 diverse provenances of ornamental rape samples, as demonstrated by the electrophoresis results (using the UBC850 marker) presented in Figure 2. Each primer pair generated a range of 4 to 5 fragments. In total, 1294 clear bands were amplified, with 1142 of these bands being polymorphic. The high level of polymorphism, at 88.3%, underscores the genetic variability among the 35 individuals. This genetic diversity is a valuable asset for assessing the genetic diversity within ornamental rape germplasm resources.

Distance is a measure of similarity between samples. The larger the genetic distance value is, the more significant the difference between varieties is. As depicted in Table S2, the genetic distances among all varieties indicate a range of genetic differentiation. The values of Nei’s genetic distance (GD) extend from a minimum of 0.066 to a maximum of 0.854. The most considerable genetic gap is observed between variety No. 4 and No. 29, with a GD of 0.854, denoting a significant genetic disparity. Conversely, the pairs with the shortest genetic distances, suggesting a high level of genetic similarity, are between No. 6 and No. 23, No. 21, No. 12 and No. 26, and No. 24 and No. 27, all of which have a GD of 0.066. In addition to distance, the similarity measure between samples also has genetic consistency. In general, the greater the genetic consistency, the closer the object. The genetic consistency ranged from 0.426 to 0.936. Therefore, at the DNA level, ornamental rape has rich genetic diversity.

As can be seen from Figure 3, the genetic similarity coefficients of the 35 ornamental rape germplasm samples range from 0.57 to 0.94. When the similarity coefficient is set at 0.65, the 35 samples are classified into 7 distinct clusters. The first category is distinguished by the unique petal color of Pink Purple No. 29, which led to its separate classification. The second group comprises Germplasm No. 4, notable for its exceptional performance in corolla width, pod length, and 1000-grain weight. The third cluster includes varieties 17, 25, and 16, which excelled in plant height, leaf length, corolla width, and 1000-grain weight, marking them as prime candidates for superior ornamental germplasm. The fourth category features White Flower No. 2, surpassing other germplasms in all nine agronomic traits evaluated. The fifth category encompasses varieties 31, 28, and 7, recognized for their impressive leaf length and the number of branches on the main stem. The sixth category is more diverse, containing 11 varieties, while the seventh comprises 15. The germplasms in these last two groups can largely be further categorized into numerous subclasses based on their flower color. The results indicate: (1) Samples 1 and 18, as well as 10 and 11, have a close genetic relationship, with a genetic similarity coefficient of 0.92, and they exhibit consistent flowering colors in terms of morphological traits; (2) Samples 6 and 21 show a high degree of similarity in flower color, with a genetic similarity coefficient reaching 1.00, suggesting that they are very likely to be the same variety; (3) Samples 31, 32, and 33, all originating from Jiangxi, China, have relatively large genetic distances from each other, indicating that the genetic diversity of ornamental rape germplasm from the same region is also rich; (4) The other samples have more distant genetic relationships. Genetic differences among the resources are evident at the molecular level, and ISSR molecular marker technology provides a clear means to differentiate the ornamental rape germplasm resources effectively.

## Discussion

As an ornamental plant, rape has seen rapid development and a surge in economic value, greatly benefiting rape growers [25]. To mitigate the unpredictable variability among germplasm resources that can be attributed to environmental factors, we cultivated 35 ornamental rape plants on a single plot. The findings revealed a coefficient of variation in agronomic traits ranging from 9.24% to 83%. Notably, even plants from the same region exhibited significant molecular-level differences, highlighting the rich genetic diversity within the species’ agronomic traits.

The genus encompasses a multitude of ornamental rape varieties that are widely distributed, exhibit high phenotypic similarity, and have ambiguous species classifications, making the identification and utilization of these germplasm resources challenging. Current research on ornamental rape is primarily concentrated on genetic levels, cultivation practices, chemical regulation, flower color, and flowering periods. Xiao et al. have demonstrated that flowering can be manipulated by factors such as temperature, light, water and soil conditions, and cultivation methods [26]. Ding et al. identified that the BNPLP1 gene influences flowering time, plant height, and the length of the main inflorescence [27]. Zhou et al. investigated SNP loci and potential candidate genes in inbred lines related to the initial, mature, and final flowering times across two environments in a panel of 300 European rape (*Brassica napus* L.) samples, uncovering 57 Brassica napus genes homologous to 39 flowering time genes in Arabidopsis [28]. In this study, 35 ornamental rape (*Brassica napus* L.) germplasm resources were analyzed for genetic diversity based on key agronomic traits and ISSR molecular markers, elucidating the genetic background and relationships of these resources. The process also involved the selection of superior ornamental rape resources, providing theoretical guidance for the further innovation and heterosis utilization in ornamental rape germplasm.

Molecular markers, as potent tools for germplasm identification and genetic diversity analysis, have been extensively employed in genetic background analysis, germplasm recovery, and the selection of breeding parents [29]. Various molecular markers have been utilized in studies to discern genetic diversity in rape (*Brassica napus* L.). Employing RAPD and EST-SSR techniques, Li et al. examined the genetic diversity among 92 cultivars from China, the USA, Canada, and select European countries, categorizing them into three groups with considerable genetic variation [30]. Furthermore, the clustering of rape plants from diverse regions underscored the inevitable impact of the ecological environment on genetic variation. Fazeli et al. utilized RAPD markers to assess genetic relationships among rapeseed genotypes from several countries, including France, Canada, Germany, Iran, Hungary, Denmark, Australia, and the United States. Cluster analysis divided the genotypes into five primary clusters, revealing genetic disparities among genotypes of the same geographical origin. This indicates that geographical origin alone cannot be the basis for achieving high heterosis in hybrid parents and necessitates precise genetic studies. The study concluded that RAPD is a straightforward, cost-effective, and expeditious method for evaluating the genetic diversity of rape [31].

ISSR, a straightforward and stable molecular marker technique, tags amplification repeat regions. High proportions of repetitive sequences in plant genomes can lead to high polymorphism rates, effectively revealing species polymorphism [32]. ISSR has been extensively applied in plant genetic diversity studies. Yang et al., using ISSR markers, uncovered the causes of population decline in sea anemones, proposed conservation strategies, and laid the scientific groundwork for the development of superior seed resources [33]. Peng et al. explored the agronomic traits, active component content, and genetic diversity of various Bupleurum chinense germplasm resources. Cluster analysis based on genetic markers grouped 13 species of Bupleurum chinense into four categories, indicating that component content does not necessarily correlate with germplasm and is susceptible to environmental influences [34]. Mesfer ALshamrani et al. investigated the genetic diversity of six sesame landraces using ISSR markers, amplifying 233 alleles with an average polymorphism percentage of 65.32%, demonstrating the high reproducibility of the ISSR markers used [35]. While ISSR molecular markers are predominantly employed to study rape diversity, few studies have focused on ornamental rape. In this study, eight primers were selected to amplify polymorphic bands, with the optimal primer being UBC850. The analysis of molecular markers and agronomic data demonstrated that ISSR markers could accurately and reliably identify genetic diversity among 35 ornamental rape cultivars and germplasm resources, underscoring their significance in the utilization, breeding, and conservation of ornamental rape germplasm.

## Acknowledgment

I extend my sincere gratitude to every author who participated in the paper; it is with their help that I have been able to complete this article.

## Supporting information

**Table S1**. Descriptive results of agronomic traits of different ornamental rape varieties

**Table S2**. Nei’s Original Measures of Genetic Identity (above diagonal) and Genetic distance (below diagonal)

**Figure 1**. Cluster maps of agronomic traits of 35 ornamental rape germplasm resources

**Figure.2**. Agarose gel electrophoresis of UBC850 ISSR primer

**Figure 3**. Cluster diagram of ornamental rape germplasm resources based on ISSR markers

## Author Contributions

Conceptualization: Xigang Dai, Mang Xia

Data curation: Mang Xia

Formal analysis: Mang Xia, Meizhu Chen,Xiaoxiao Dong Investigation: Mang Xia, Jingdong Chen, Miao Cheng

Methodology: Xigang Dai, Mang Xia

Project administration: Xigang Dai

Writing – original draft: Mang Xia

Writing – review & editing: Heping Wan, Yuanhuo Dong, Changli Zeng and Xigang Dai

## Funding

This research was funded by the Key research and development project of Hubei Province: 2022BBA0064; the National science and technology support program of China (2022ZD04010), and Jianghan University scientific research project funding scheme (2022XKZX17).

## Notes

### Competing Interest Statement

The authors have declared no competing interest.

